# The repurposed STAT3 inhibitor pyrimethamine controls mycobacterial infection-induced vascular permeability and mycobacterial burden

**DOI:** 10.1101/2024.11.28.625803

**Authors:** Darryl JY Han, Xiangyu Sui, Ria Sorayah, Luke H Hoeppner, Stefan H Oehlers

## Abstract

Infection-induced vascular pathologies are a side effect of the immune response to contact with a range of pathogens. Mycobacteria, including *Mycobacterium tuberculosis*, are particularly adept at co-opting vascular leakiness as a survival mechanism to shape the host immune response and impede the delivery of antibiotics to sites of infection. Here using the zebrafish-*Mycobacterium marinum* infection model, we confirm a critical role for Signal transducer and activator of transcription 3 (STAT3) in mediating infection-induced vascular permeability, and demonstrate the ability of FDA-approved drugs atovaquone and pyrimethamine to restore vascular barrier function without compromising immune control of mycobacterial infection. Additionally, we find an antibiotic effect of pyrimethamine against *Mycobacterium marinum* via inhibition of bacterial dihydrofolate reductase. Together our findings suggest pyrimethamine could be used as adjunctive therapy against mycobacterial infection.

## Main text

Active tuberculosis is associated with host vascular pathologies including angiogenesis and vascular permeability driven by the production of vasoactive molecules such as Vascular endothelial growth factor (VEGF) and Angiopoietin-2 by infected macrophages ^1-5^. Infection-induced angiogenesis serves to supply granuloma-bound mycobacteria with oxygen, delaying hypoxia-induced growth arrest ^3^. Both the new blood vessels and nearby existing vasculature are permeable allowing the leak of contents into the surrounding tissue and egress of neutrophils while preventing efficient delivery of antibiotics and T cells ^6, 7^, which can aggravate late stage tuberculosis granulomas. Current options for vascular normalizing adjunctive therapy include VEGF pathway targeting drugs and biologics developed for cancer that are prohibitively expensive for mass deployment ^3, 7, 8^.

The pleiotropic molecule Signal transducer and activator of transcription 3 (STAT3) has been shown to be mediate the transduction of VEGF-induced vascular leakiness in endothelial cells across zebrafish, mice, and humans ^9^. Targeting STAT3 has also shown promise as a host-directed therapy in preclinical tuberculosis models suggesting that inhibition may not be detrimental to the immune control of mycobacterial infection ^10-12^. We first utilised genetic tools to knockdown *stat3* expression with CRISPR-Cas9 gene editing creating F0 “crispants” with reduced transcription of *stat3* measured by RT-qPCR (Supplementary Figure 1). We then sought to determine if Stat3 controlled infection-induced vascular permeability by infecting mosaic *stat3* crispant zebrafish embryos with *M. marinum* and measuring vascular permeability around granulomas (Figure 1A). Depletion of Stat3 did not affect bacterial burden (Figure 1B), but did decrease vascular leakiness (Figure 1C). These results were confirmed in experiments with the *stat3*^*zf3708*^ allele (Figure 1D-E). Together these data demonstrate the feasibility of targeting STAT3 to reduce infection-induced vascular pathology.

**Figure 1:**
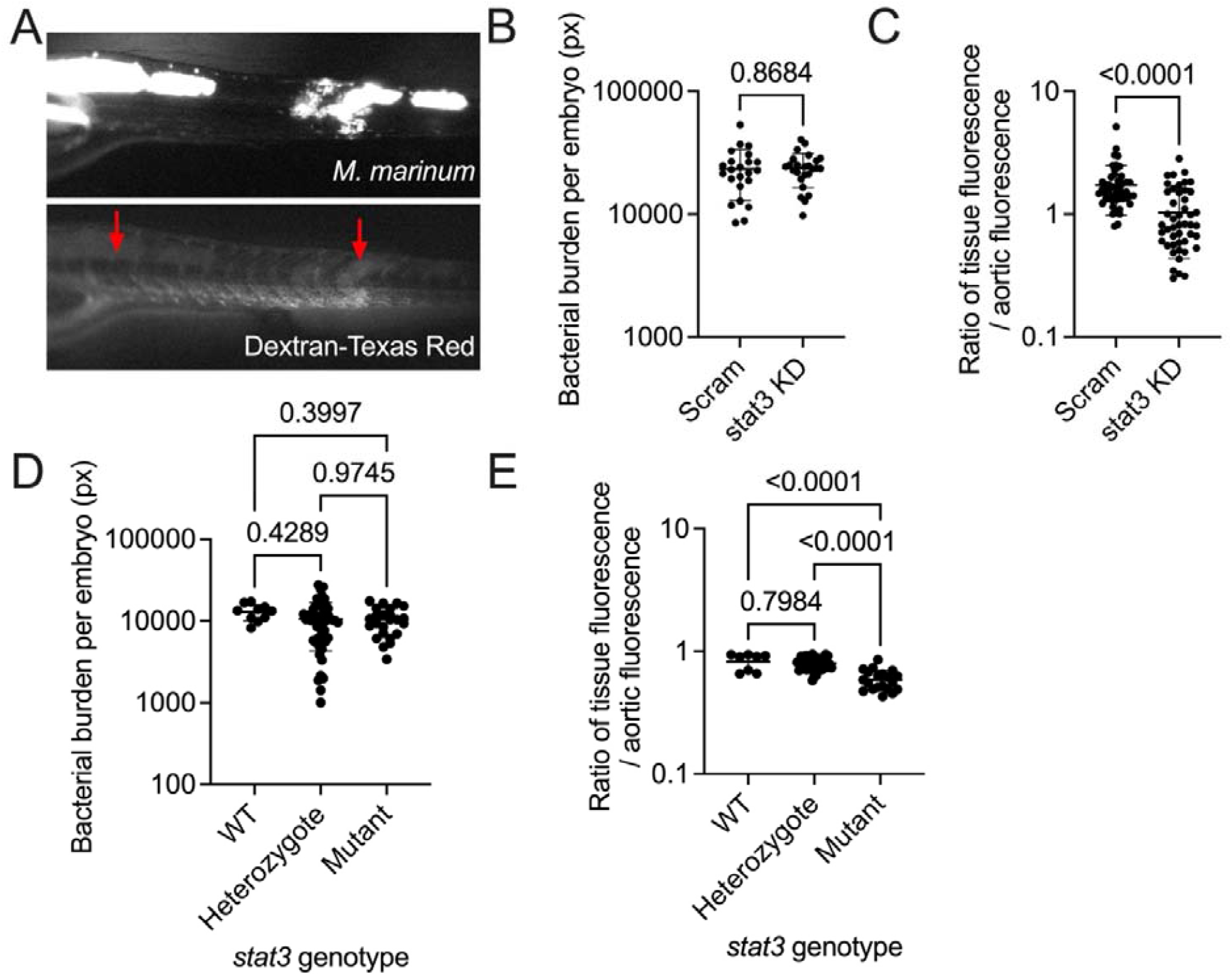
Genetic depletion of STAT3 reduces infection-induced vascular permeability (A) Microangiography of 6 dpi *M. marinum*-infected zebrafish embryos allows quantification of vascular permeability by fluorescent microscopy. Red arrows indicate areas of infection-induced vascular leak. (B) Quantification of *M. marinum* burden in 6 dpi trunk infected *stat3* crispant zebrafish embryos. (C) Quantification of infection-induced vascular permeability in 6 dpi trunk infected *stat3* crispant zebrafish embryos. (D) Quantification of *M. marinum* burden in 6 dpi trunk infected *stat3* mutant zebrafish embryos. (E) Quantification of infection-induced vascular permeability in 6 dpi trunk infected *stat3* mutant zebrafish embryos.

Two widely used FDA approved anti-parasitic drugs have been found to inhibit STAT3 activity and maybe be suitable candidates for drug repurposing due to their widespread availability and known safety profiles, atovaquone (Mepron, FDA-approved over 20 years ago) and pyrimethamine (Daraprim, WHO essential medicine) ^13, 14^. We examined the effect of atovaquone and pyrimethamine treatment on vascular permeability during established mycobacterial infection by infecting zebrafish embryos with *M. marinum* and then treating at 6 days post infection (dpi) for one day before performing the vascular permeability assay at 7 dpi (Figure 2A). This short treatment did not affect bacterial burden (Figure 2B), and significantly reduced infection-induced vascular permeability (Figure 2C).

**Figure 2:**
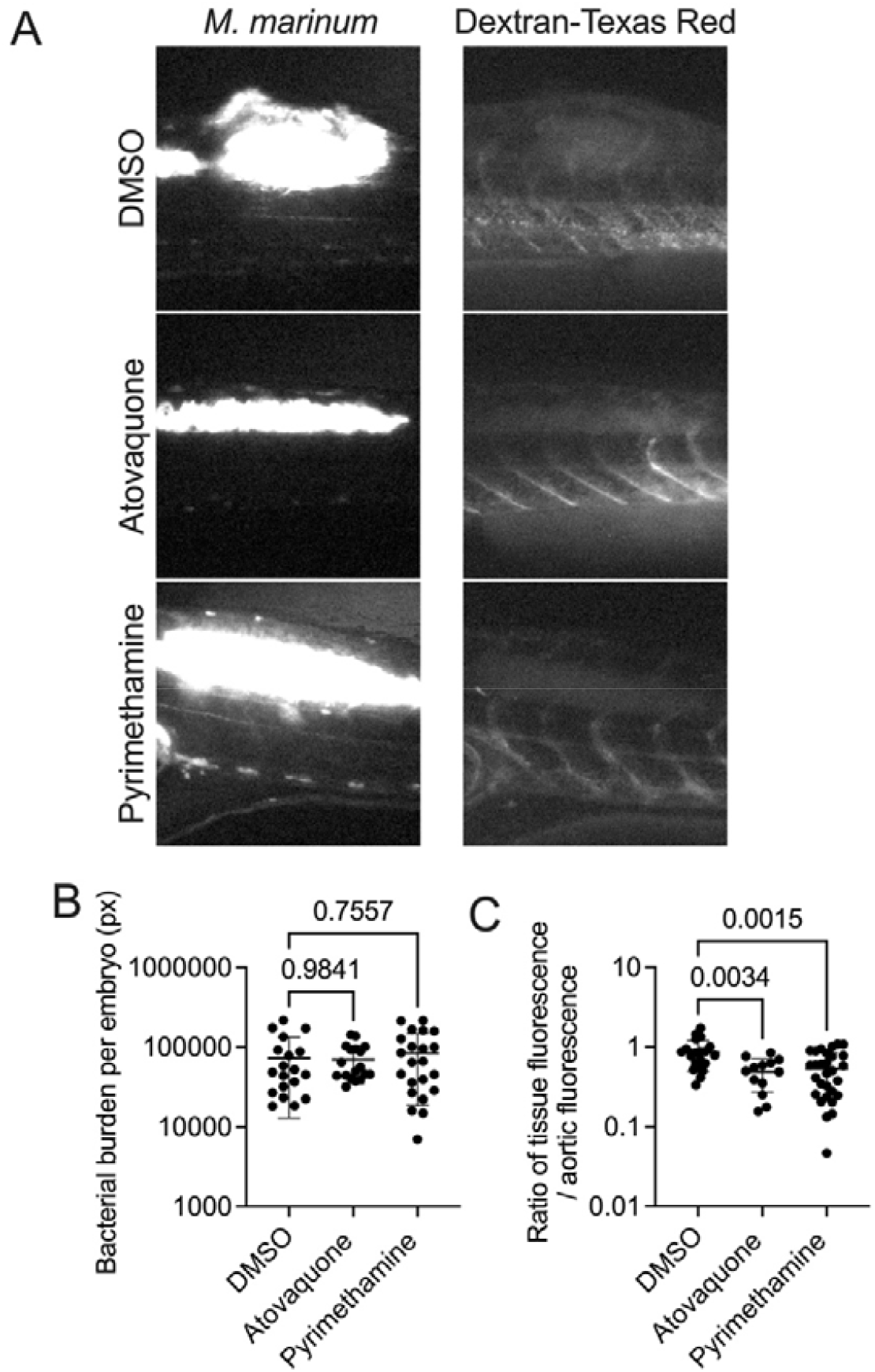
Inhibition of STAT3 with repurposed small molecules reduces infection-induced vascular permeability (A) Representative microangiography images of *M. marinum*-infected zebrafish treated with 50 µM atovaquone or pyrimethamine. (B) Quantification of *M. marinum* burden in trunk infected zebrafish embryos treated with 50 µM atovaquone or pyrimethamine from 6-7 dpi. (C) Quantification of infection-induced vascular permeability in trunk infected zebrafish embryos treated with 50 µM atovaquone or pyrimethamine from 6-7 dpi.

We next investigated if atovaquone and pyrimethamine treatment could potentiate isoniazid antibiotic efficacy using an established infection model treated with isoniazid and atovaquone or pyrimethamine at 5 dpi and assayed two days later at 7 dpi. Treatment with atovaquone and pyrimethamine did not affect the efficacy of isoniazid delivered by soaking or infusion (Figure 3A and 3B). Interestingly we observed reduced *M. marinum* burden in animals treated with pyrimethamine alone suggesting a direct antibiotic effect in addition to inhibition of host Stat3 in the zebrafish-*M. marinum* infection system (Figure 3A and 3B). We confirmed this observation in systemically infected animals treated with pyrimethamine from 0-5 dpi (Figure 3C).

**Figure 3:**
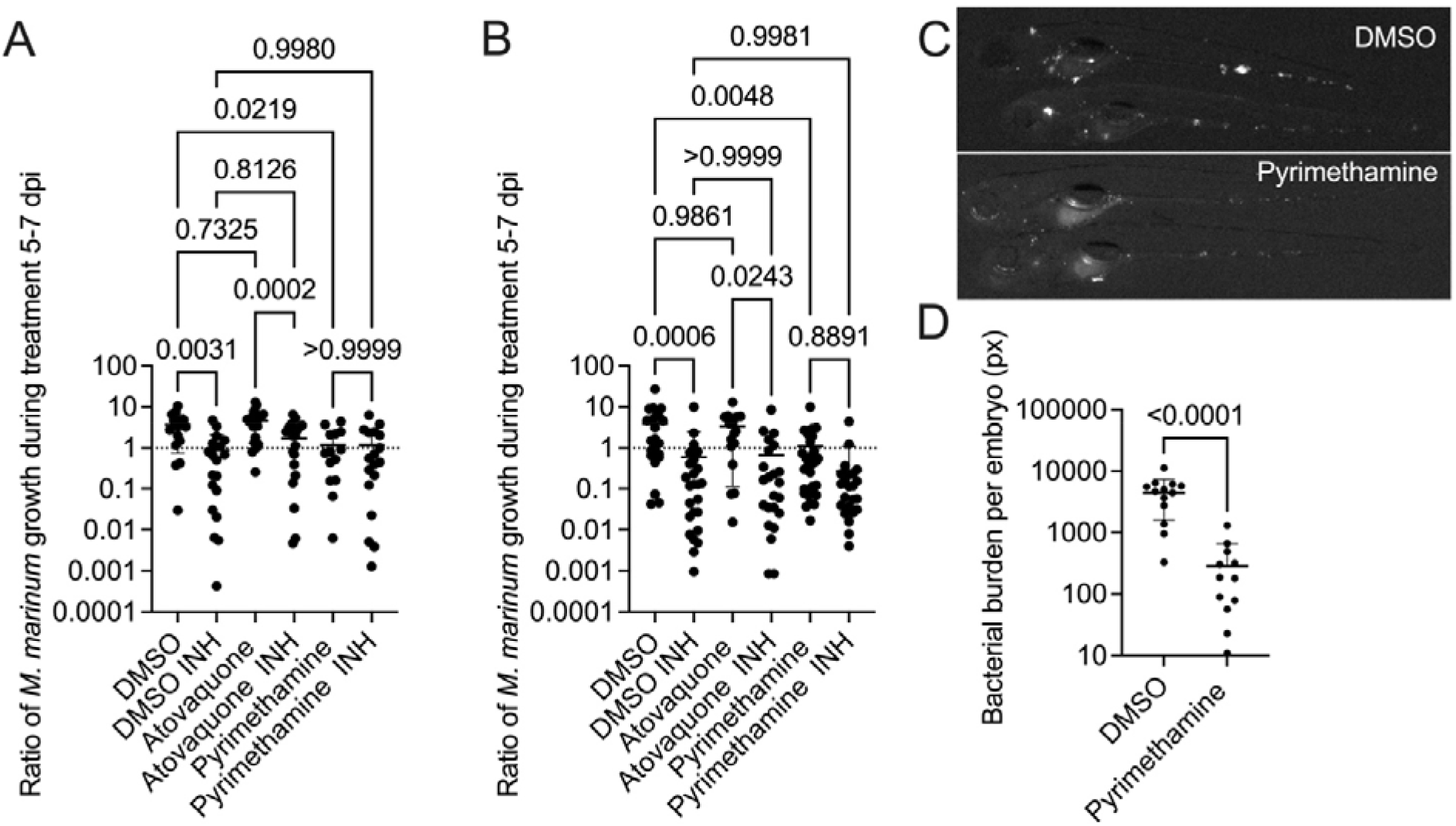
Atovaquone and pyrimethamine do not interact with isoniazid, and pyrimethamine treatment inhibits *M. marinum* growth in zebrafish embryos (A) Quantification of *M. marinum* burden in trunk infected zebrafish embryos treated with 50 µM atovaquone or pyrimethamine and 200 µM isoniazid (INH) from 5-7 dpi by immersion and infused with isoniazid (INH) at 5 dpi. (B) Quantification of *M. marinum* burden in trunk infected zebrafish embryos treated with 50 µM atovaquone or pyrimethamine from 5-7 dpi by immersion and infused with isoniazid (INH) at 5 dpi. (C) Representative images of 5 dpi *M. marinum*-infected zebrafish embryos treated with 25 µM pyrimethamine from 0-5 dpi. (D) Quantification of *M. marinum* burden in systemically infected embryos treated with 25 µM pyrimethamine from 0-5 dpi.

Pyrimethamine is an inhibitor of bacterial dihydrofolate reductase (DHFR), albeit for non-mycobacterial species, and there has been recent interest in targeting mycobacterial DHFR with anti-parasitic inhibitors ^15^. Of clinical relevance, a protective effect of prophylactic pyrimethamine prescription against tuberculosis has been observed in population level studies suggesting activity against mycobacteria ^16, 17^. While atovaquone did not show any effect on *in vitro M. marinum* growth (Figure 4A), pyrimethamine exhibited a direct antibiotic effect on *M. marinum* in broth culture (Figure 4B). No effect was seen on *M. abscessus* in broth culture suggesting specificity of activity to individual species of mycobacteria that will be important to follow up across a wider range of pathogenic species (Figure 4C). Partial rescue of growth in pyrimethamine-treated *M. marinum* broth cultures by supplementation of tetrahydrofolic acid, the metabolic product of DHFR, suggest the antibiotic effect is at least partially due to DHFR inhibition (Figure 4D).

**Figure 4:**
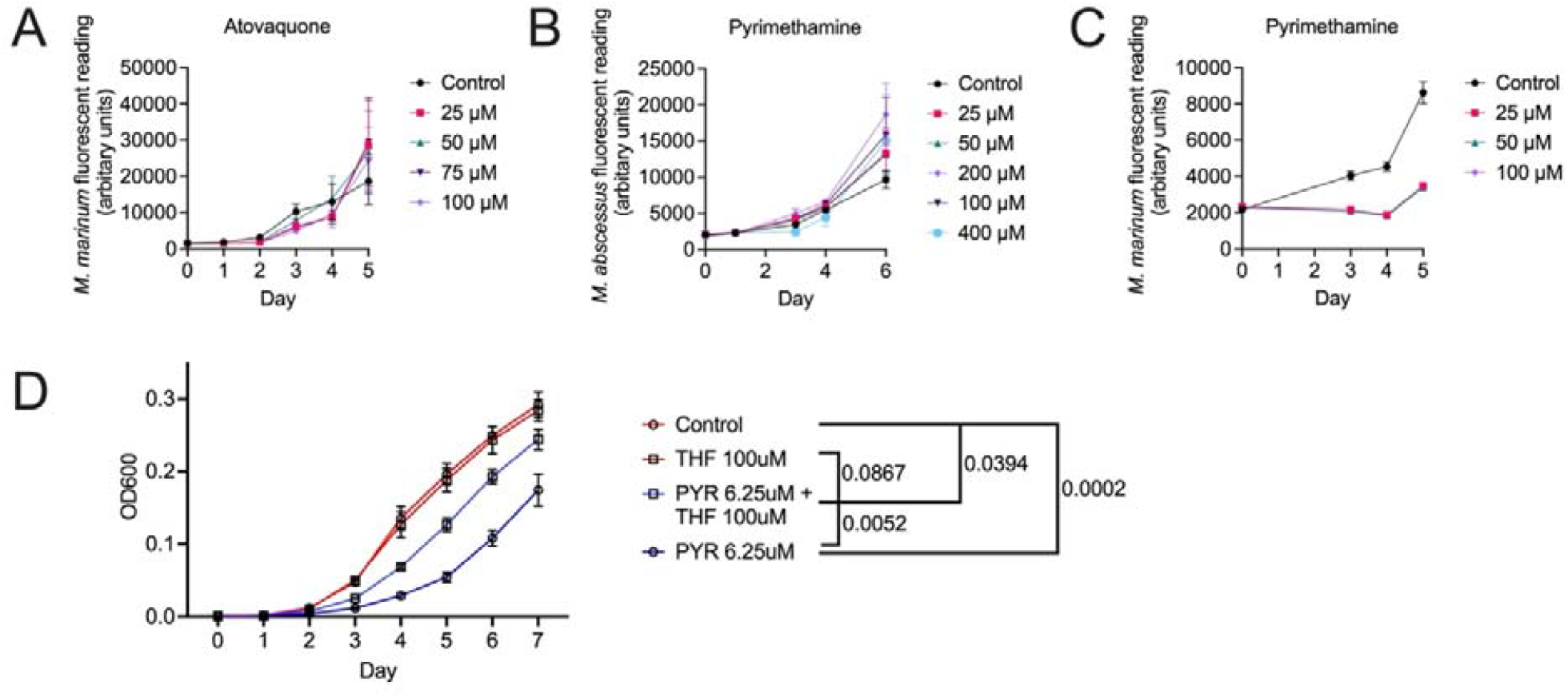
Direct antibiotic effect of pyrimethamine against *M. marinum*, but not *M. abscessus*, cultures *in vitro* (A) Growth curve of *M. marinum* treated with atovaquone as indicated. (B) Growth curve of *M. marinum* treated with pyrimethamine as indicated. (C) Growth curve of *M. abscessus* treated with pyrimethamine as indicated. (D) Growth curve of *M. marinum* treated with 6.25 μM pyrimethamine (PYR) and 100 μM tetrahydrofolic acid (THF). Statistical comparisons are of technical replicates of Day 7 OD600 readings by ANOVA.

Together, our data demonstrate the feasibility of targeting Stat3 with widely available FDA-approved anti-parasitic drugs to reduce infection-induced vascular pathology and describe the susceptibility of *M. marinum* to pyrimethamine. Atovaquone within interacts with rifampin and rifabutin, important TB antibiotics, resulting in decreased atovaquone serum concentration which may limit the efficacy of atovaquone as an adjunctive therapy ^18^. Conversely, pyrimethamine is well tolerated and is not documented to interact with frontline TB drugs making it an excellent candidate for immediate repurposing into an adjunctive host-directed therapy, or refinement of DHFR inhibitor activity, against mycobacterial infections.

## METHODS

### Bacterial culture

Fluorescent M strain *M. marinum* and *M. abscessus* ATCC19977 carrying pTEC plasmids driving expression of TdTomato or Wasabi under the control of the Mycobacterium Strong Promoter were cultured in 7H9 media supplemented with 50 μg/ml hygromycin B and single cell suspensions were prepared for injection as previously described ^19^.

Atovaquone, pyrimethamine and tetrahydrofolic acid dissolved in DMSO were added to 7H9 cultures to the final concentrations indicated. Growth curves were carried out in 200 µL of 7H9 culture in 96 well. Plates were incubated at 30°C without shaking and fluorescence was measured at each time point shown with a Tecan plate reader (excitation 554 nm, emission 581 nm).

### Zebrafish infection procedure

Animal experiments were approved by the A*STAR IACUC under protocols 211667 and 221694. Experimental embryos were performed by natural spawning of adult zebrafish and raised at 28°C in E3 embryo media supplemented with methylene blue for one day.

Embryos were dechorionated with pronase and subsequently raised in E3 supplemented with 45 µg/mL 1-phenyl 2-thiourea (PTU, Sigma-Aldrich) from one day post fertilization. Two day post fertilization embryos were anesthetized with 160 µg/mL tricaine and microinjected with ∼200 CFU *M. marinum* into the neural tube (trunk) for vascular permeability assays or the caudal plexus blood vessels for systemic infections. Infected embryos were recovered into E3 supplemented with PTU and raised at 28°C under drug treatments as indicated.

### CRISPR-Cas9 editing of *stat3*

A set of 4 *stat3*-targeting gRNAs were *in vitro* transcribed, annealed to Cas9 enzyme (), and injected into the yolks of 1-4 cell stage zebrafish embryos ^20^. Oligo sequences are in Table 1. Knockdown efficiency was verified by qRT-PCR with >50% depletion of transcript observed.

**Table 1.**
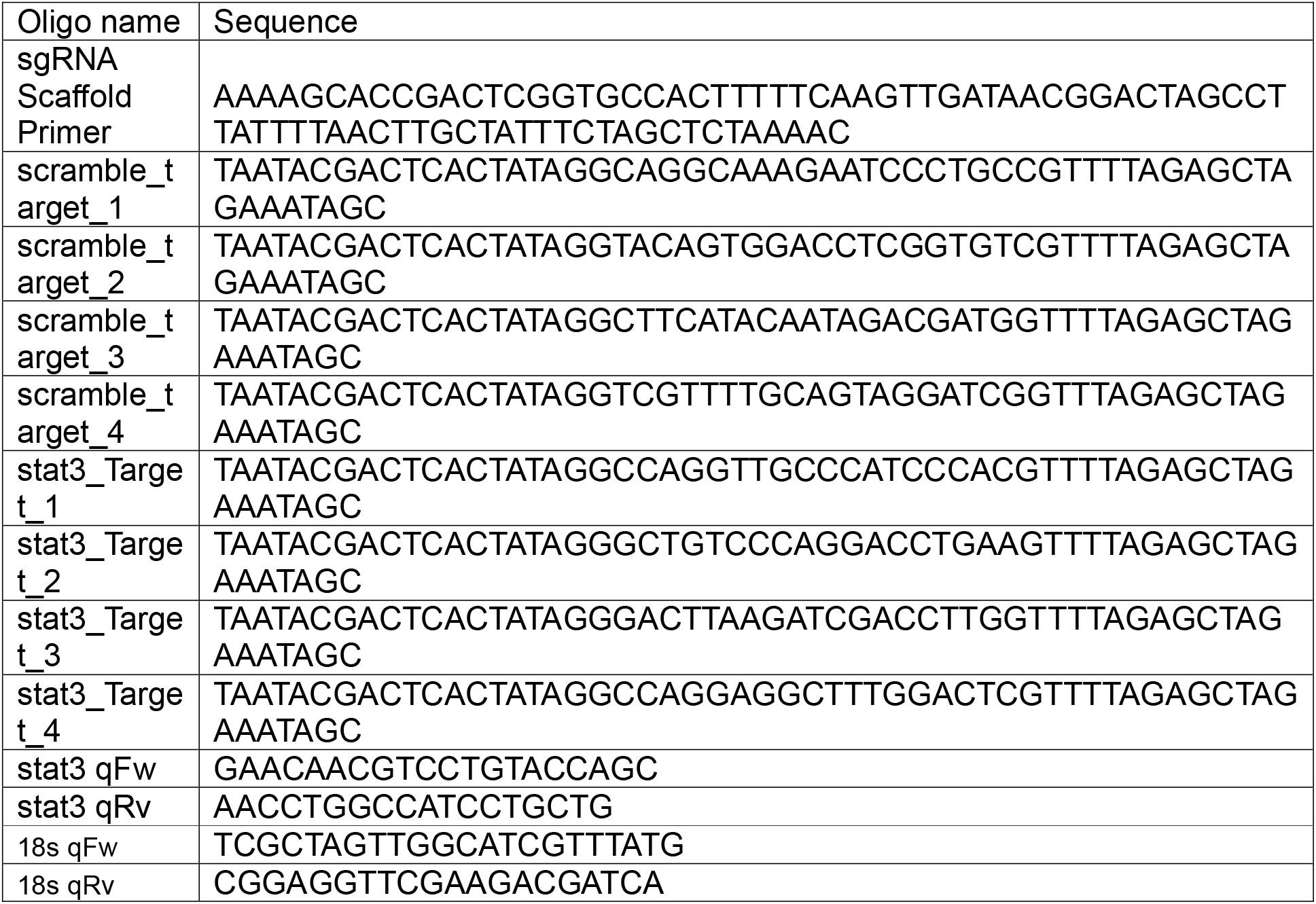
Sequences of oligonucleotides used in this study.

### Drug treatment

STAT3 inhibitors atovaquone and pyrimethamine were dissolved in DMSO, isoniazid was dissolved in water. Soaking was achieved by addition of concentrated drug directly into the E3 media of infected embryos to achieve the final concentrations as stated. Infusion of isoniazid was performed by microinjection of 10 nl of an aqueous solution of 400 µM isoniazid into the dorsal aorta or caudal vein of infected anesthetized zebrafish.

### Bacterial burden measurement

Anesthetized larvae were imaged by epifluorescence microscopy under consistent exposure and camera settings within each experiment. Bacterial fluorescent pixel count was carried out in ImageJ as previously described ^21^.

### Vascular permeability assay

Anesthetized larvae were perfused with a solution of 1 mg/ml dextran-Texas Red 70,000 MW (Thermofisher) by microinjection into the dorsal aorta or caudal vein ^2^. Injected larvae were recovered to E3 for 30 minutes to facilitate circulation of the dye prior to being re-anesthetized and imaged. Vascular leakage was calculated as the ratio of the maximum Texas Red intensity in the granuloma divided by the aorta ventral to the granuloma.

### Statistical analyses

All data are presented as mean +/-standard deviation. Statistical comparisons were performed by *t* tests for two way comparisons or ANOVA for experiments with three or more groups.

## Author Contributions

DJYH performed experiments, analyzed data, prepared display items. XS performed experiments, analyzed data.

RS performed experiments, analyzed data. LHH conceived study and provided reagents.

SHO conceived study, performed experiments, analyzed data, wrote manuscript draft. All authors reviewed the manuscript.

## Notes

No competing financial interests have been declared.

## ACKNOWLEDGMENT

IMCB Zebrafish Facility for expert management of our breeding stock. Dr Bruno Reversade for use of his microscopy room and equipment.

## REFERENCES

(1) Datta, M.; Via, L. E.; Kamoun, W. S.; Liu, C.; Chen, W.; Seano, G.; Weiner, D. M.; Schimel, D.; England, K.; Martin, J. D.; et al. Anti-vascular endothelial growth factor reatment normalizes tuberculosis granuloma vasculature and improves small molecule delivery. Proc Natl Acad Sci U S A 2015, 112 (6), 1827–1832. DOI: 10.1073/pnas.1424563112.

(2) Oehlers, S. H.; Cronan, M. R.; Beerman, R. W.; Johnson, M. G.; Huang, J.; Kontos, C. D.; Stout, J. E.; Tobin, D. M. Infection-induced vascular permeability aids mycobacterial growth. J Infect Dis 2017, 215 (5), 813–817. DOI: 10.1093/infdis/jiw355.

(3) Oehlers, S. H.; Cronan, M. R.; Scott, N. R.; Thomas, M. I.; Okuda, K. S.; Walton, E. M.; Beerman, R. W.; Crosier, P. S.; Tobin, D. M. Interception of host angiogenic signalling limits mycobacterial growth. Nature 2015, 517 (7536), 612–615. DOI: 10.1038/nature13967.

(4) Walton, E. M.; Cronan, M. R.; Cambier, C. J.; Rossi, A.; Marass, M.; Foglia, M. D.; Brewer, W. J.; Poss, K. D.; Stainier, D. Y. R.; Bertozzi, C. R.; et al. Cyclopropane Modification of Trehalose Dimycolate Drives Granuloma Angiogenesis and Mycobacterial Growth through Vegf Signaling. Cell Host Microbe 2018, 24 (4), 514–525 e516. DOI: 10.1016/j.chom.2018.09.004.

(5) Brewer, W. J.; Xet-Mull, A. M.; Yu, A.; Sweeney, M. I.; Walton, E. M.; Tobin, D. M. Macrophage NFATC2 mediates angiogenic signaling during mycobacterial infection. Cell reports 2022, 41 (11), 111817. DOI: 10.1016/j.celrep.2022.111817 From NLM Medline.

(6) Kam, J. Y.; Cheng, T.; Garland, D. C.; Britton, W. J.; Tobin, D. M.; Oehlers, S. H. Inhibition of infection-induced vascular permeability modulates host leukocyte recruitment to Mycobacterium marinum granulomas in zebrafish. Pathogens and disease 2022, 80 (1). DOI: 10.1093/femspd/ftac009.

(7) Datta, M.; Via, L. E.; Dartois, V.; Weiner, D. M.; Zimmerman, M.; Kaya, F.; Walker, A. M.; Fleegle, J. D.; Raplee, I. D.; McNinch, C.; et al. Normalizing granuloma vasculature and matrix improves drug delivery and reduces bacterial burden in tuberculosis-infected rabbits. Proc Natl Acad Sci U S A 2024, 121 (14), e2321336121. DOI: 10.1073/pnas.2321336121 From NLM Medline.

(8) Polena, H.; Boudou, F.; Tilleul, S.; Dubois-Colas, N.; Lecointe, C.; Rakotosamimanana, N.; Pelizzola, M.; Andriamandimby, S. F.; Raharimanga, V.; Charles, P.; et al. Mycobacterium tuberculosis exploits the formation of new blood vessels for its dissemination. Scientific reports 2016, 6, 33162. DOI: 10.1038/srep33162.

(9) Wang, L.; Astone, M.; Alam, S. K.; Zhu, Z.; Pei, W.; Frank, D. A.; Burgess, S. M.; Hoeppner, L. H. Suppressing STAT3 activity protects the endothelial barrier from VEGF-mediated vascular permeability. Dis Model Mech 2021, 14 (11). DOI: 10.1242/dmm.049029.

(10) Upadhyay, R.; Sanchez-Hidalgo, A.; Wilusz, C. J.; Lenaerts, A. J.; Arab, J.; Yeh, J.; Stefanisko, K.; Tarasova, N. I.; Gonzalez-Juarrero, M. Host Directed Therapy for Chronic Tuberculosis via Intrapulmonary Delivery of Aerosolized Peptide Inhibitors Targeting the IL-10-STAT3 Pathway. Scientific reports 2018, 8 (1), 16610. DOI: 10.1038/s41598-018-35023-0 From NLM Medline.

(11) Lastrucci, C.; Benard, A.; Balboa, L.; Pingris, K.; Souriant, S.; Poincloux, R.; Al Saati, T.; Rasolofo, V.; Gonzalez-Montaner, P.; Inwentarz, S.; et al. Tuberculosis is associated with expansion of a motile, permissive and immunomodulatory CD16(+) monocyte population via the IL-10/STAT3 axis. Cell Res 2015, 25 (12), 1333–1351. DOI: 10.1038/cr.2015.123 From NLM Medline.

(12) Gao, Y.; Basile, J. I.; Classon, C.; Gavier-Widen, D.; Yoshimura, A.; Carow, B.; Rottenberg, M. E. STAT3 expression by myeloid cells is detrimental for the T-cell-mediated control of infection with Mycobacterium tuberculosis. PLoS Pathog 2018, 14 (1), e1006809. DOI: 10.1371/journal.ppat.1006809 From NLM Medline.

(13) Xiang, M.; Kim, H.; Ho, V. T.; Walker, S. R.; Bar-Natan, M.; Anahtar, M.; Liu, S.; Toniolo, P. A.; Kroll, Y.; Jones, N.; et al. Gene expression-based discovery of atovaquone as a STAT3 inhibitor and anticancer agent. Blood 2016, 128 (14), 1845–1853. DOI: 10.1182/blood-2015-07-660506 From NLM Medline.

(14) Khan, M. W.; Saadalla, A.; Ewida, A. H.; Al-Katranji, K.; Al-Saoudi, G.; Giaccone, Z. T.; Gounari, F.; Zhang, M.; Frank, D. A.; Khazaie, K. The STAT3 inhibitor pyrimethamine displays anti-cancer and immune stimulatory effects in murine models of breast cancer. Cancer Immunol Immunother 2018, 67 (1), 13–23. DOI: 10.1007/s00262-017-2057-0 From NLM Medline.

(15) Andrade Meirelles, M.; Almeida, V. M.; Sullivan, J. R.; de Toledo, I.; Dos Reis, C. V.; Cunha, M. R.; Zigweid, R.; Shim, A.; Sankaran, B.; Woodward, E. L.; et al. Rational Exploration of 2,4-Diaminopyrimidines as DHFR Inhibitors Active against Mycobacterium abscessus and Mycobacterium avium, Two Emerging Human Pathogens. J Med Chem 2024. DOI: 10.1021/acs.jmedchem.4c01594 From NLM Publisher.

(16) Haller, L.; Sossouhounto, R.; Coulibaly, I. M.; Dosso, M.; Kone, M.; Adom, H.; Meyer, U. A.; Betschart, B.; Wenk, M.; Haefeli, W. E.; et al. Isoniazid plus sulphadoxine-pyrimethamine can reduce morbidity of HIV-positive patients treated for tuberculosis in Africa: a controlled clinical trial. Chemotherapy 1999, 45 (6), 452–465. DOI: 10.1159/000007239 From NLM Medline.

(17) Opravil, M.; Pechere, M.; Lazzarin, A.; Heald, A.; Ruttimann, S.; Iten, A.; Furrer, H.; Oertle, D.; Praz, G.; Vuitton, D. A.; et al. Dapsone/pyrimethamine may prevent mycobacterial disease in immunosuppressed patients infected with the human immunodeficiency virus. Clin Infect Dis 1995, 20 (2), 244–249. DOI: 10.1093/clinids/20.2.244 From NLM Medline.

(18) Sousa, M.; Pozniak, A.; Boffito, M. Pharmacokinetics and pharmacodynamics of drug interactions involving rifampicin, rifabutin and antimalarial drugs. J Antimicrob Chemother 2008, 62 (5), 872–878. DOI: 10.1093/jac/dkn330 From NLM Medline.

(19) Takaki, K.; Davis, J. M.; Winglee, K.; Ramakrishnan, L. Evaluation of the pathogenesis and treatment of Mycobacterium marinum infection in zebrafish. Nat Protoc 2013, 8 (6), 1114–1124. DOI: 10.1038/nprot.2013.068.

(20) Wu, R. S.; Lam, II; Clay, H.; Duong, D. N.; Deo, R. C.; Coughlin, S. R. A Rapid Method for Directed Gene Knockout for Screening in G0 Zebrafish. Dev Cell 2018, 46 (1), 112–125 e114. DOI: 10.1016/j.devcel.2018.06.003.

(21) Matty, M. A.; Oehlers, S. H.; Tobin, D. M. Live Imaging of Host-Pathogen Interactions in Zebrafish Larvae. Methods Mol Biol 2016, 1451, 207–223. DOI: 10.1007/978-1-4939-3771-4_14.

